# Confirmed Occurrence and Morphological Documentation of Stenopus spinosus (Risso, 1827), from Tenerife, Canary Islands

**DOI:** 10.1101/2025.08.07.669097

**Authors:** Michael Bommerer

## Abstract

This dataset provides photographic and morphological documentation of *Stenopus spinosus* Risso, 1827, based on a single adult specimen collected from Granadilla, Tenerife, Canary Islands, on 6 August 2025. The individual was observed and captured at a depth of 6 meters in a rocky overhead marine environment during a night dive, and subsequently documented through standardized dorsal-view laboratory imaging (Figures C–1 and D–1). Morphometric data were recorded, including carapace length (CL), rostrum length (RL), chela length and cheliped length. This record contributes to the regional documentation of marine decapods in the eastern Atlantic and supports ongoing biodiversity assessment efforts in the Canary Islands. Additionally, a separate in situ underwater observation of S. spinosus, recorded during a night dive in Jaca, Tenerife on 1 November 2024, is cited for comparative purposes.

## 1. Introduction

*Stenopus spinosus* Risso, 1827 is a marine decapod crustacean belonging to the family *Stenopodidae*, within the infraorder *Stenopodidea*. The species was first described by Antoine Risso (1826–1827) in his seminal work *Histoire naturelle des principales productions de l’Europe méridionale*, based on specimens from the Mediterranean coast near Nice. Since its original description, *S. spinosus* has been documented across a wide geographic range, including the eastern Atlantic Ocean, Mediterranean Sea, Gulf of Mexico, Caribbean Sea, and the Red Sea.

Bathymetric records confirm its occurrence from shallow waters to moderately deep waters, with depth records ranging from approximately 10 to 190 meters (DecaNet eds. 2025; WoRMS 2025). It is most commonly associated with rocky substrates, coral reefs, and marine caves, where it resides under ledges or within crevices during the day and emerges nocturnally to forage. Regional records from Europe include the Adriatic Sea, Aegean Sea, Balearic Sea, Alboran Sea, Ionian Sea, and Levantine Basin, as well as Atlantic coastal zones off Spain, Greece, Cameroon, and the Republic of the Congo.

Although *S. spinosus* is considered relatively common in portions of its range, published photographic and morphometric documentation from the Macaronesian region—including the Canary Islands—remains limited. The current dataset addresses this gap by providing verifiable imagery and morphological data for a single adult specimen collected from a rocky overhead environment at 6 meters depth off Granadilla, Tenerife (Canary Islands) on 6 August 2025. The specimen was documented using standardized laboratory photography and measured using decapod morphological protocols, including carapace length (CL), rostrum length (RL), and cheliped length.

Additionally, an underwater observation of a conspecific individual from Jaca, Tenerife (28°07′00″ N, 16°27′45″ W) recorded on 1 November 2024 during a night dive is included to provide ecological context and support visual comparisons. Together, these records contribute to the growing documentation of marine crustacean biodiversity in the northeastern Atlantic and serve as reference material for future faunal and ecological assessments involving *Stenopus spp*. in the Canary Islands and surrounding regions.

## 2. Materials and Methods

### 2.1 Specimen Collection and Habitat

An adult specimen of *Stenopus spinosus* was collected on 6 August 2025 during a recreational night dive at 01:00 local time off Granadilla, Tenerife, Canary Islands (28°05’07”N 16°29’23”W). The specimen was encountered at a depth of approximately 6 meters within a rocky overhead environment, where it was observed resting near the entrance of a small cave. No additional individuals were recorded in the vicinity at the time of collection. Environmental parameters such as water temperature and salinity were not recorded.

### 2.2 Photography

The specimen was transferred to a laboratory facility and photographed in dorsal view under controlled lighting conditions using a high-resolution digital camera (Canon 80D). A millimeter scale bar was included in each image for size reference. Two dorsal-view images (Figures C–1 and D–1) were captured to document the general morphology. Image contrast and sharpness were adjusted using Adobe software to improve visual clarity without altering morphological details.

**Figure C–1.**
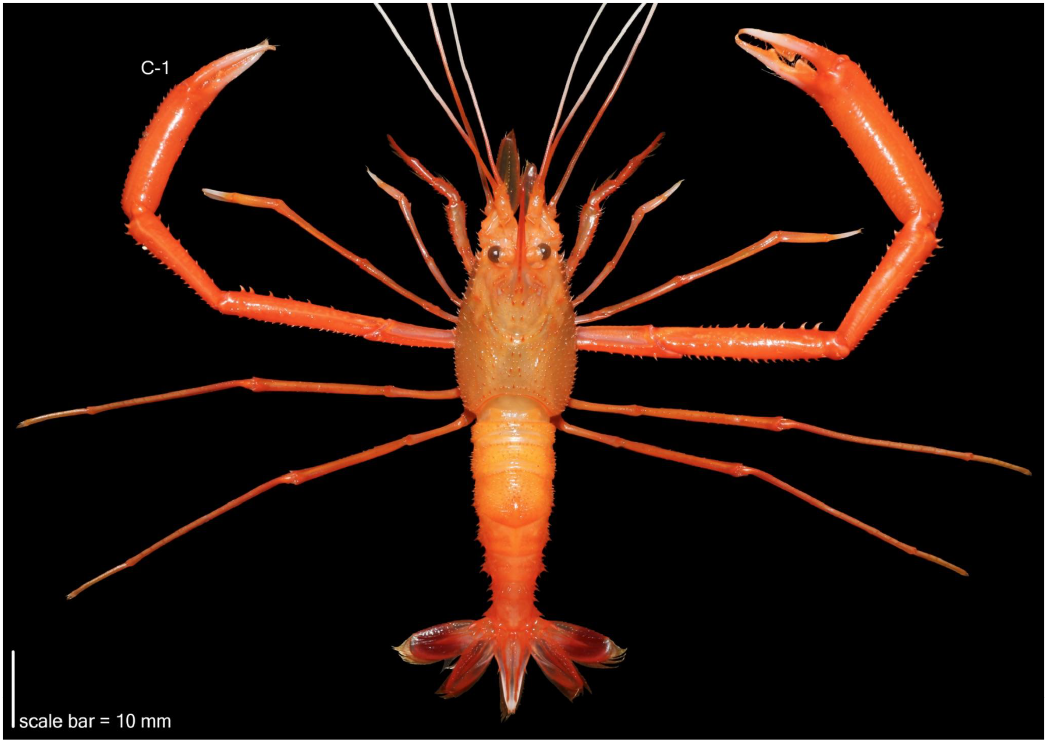
Dorsal view of *Stenopus spinosus* Risso, 1827 (Decapoda: *Stenopodidea*: *Stenopodidae*), adult specimen (Specimen ID: IMS-SSP-001) collected from Granadilla, Tenerife, Canary Islands (28°05′07″N, 16°29′23″W). The individual was encountered at a depth of 6 meters in a rocky overhead marine environment during a night dive at approximately 01:00 local time on 6 August 2025. Photographed under standardized laboratory conditions. Morphometric measurements: carapace length (CL) = 16.4 mm; rostrum length (RL) = 11.1 mm; right chela length = 36.7 mm; right cheliped length = 94.3 mm. Scale bar = 10 mm. Associated iNaturalist record: https://www.inaturalist.org/observations/304303255

**Figure D–1.**
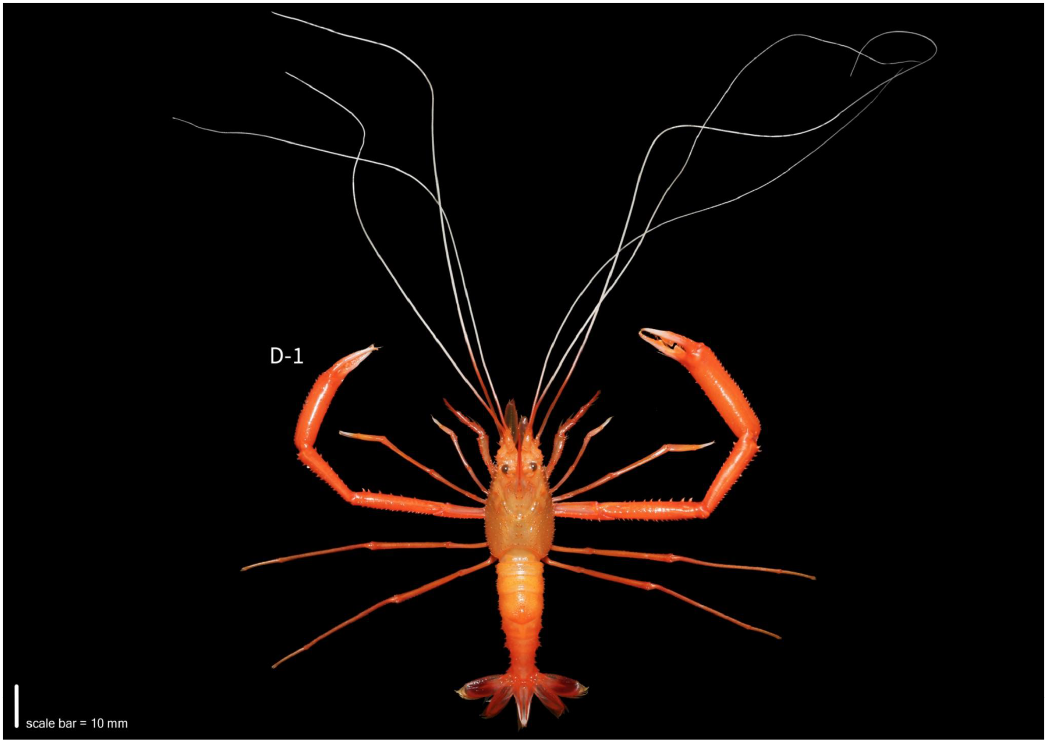
Complete dorsal view of Stenopus spinosus Risso, 1827 (Decapoda: Stenopodidea: Stenopodidae), adult specimen (Specimen ID: IMS-SSP-001) showing extended antennal appendages. Collected at a depth of 6 meters within a rocky overhead marine environment during a night dive (approx. 01:00 local time) on 6 August 2025. Locality: Granadilla, Tenerife, Canary Islands (28°05′07″N, 16°29′23″W). Photographed under standardized laboratory conditions. Morphometric measurements: carapace length (CL) = 16.4 mm; rostrum length (RL) = 11.1 mm; right chela length = 36.7 mm. Scale bar = 10 mm.

**Figure F–1.**
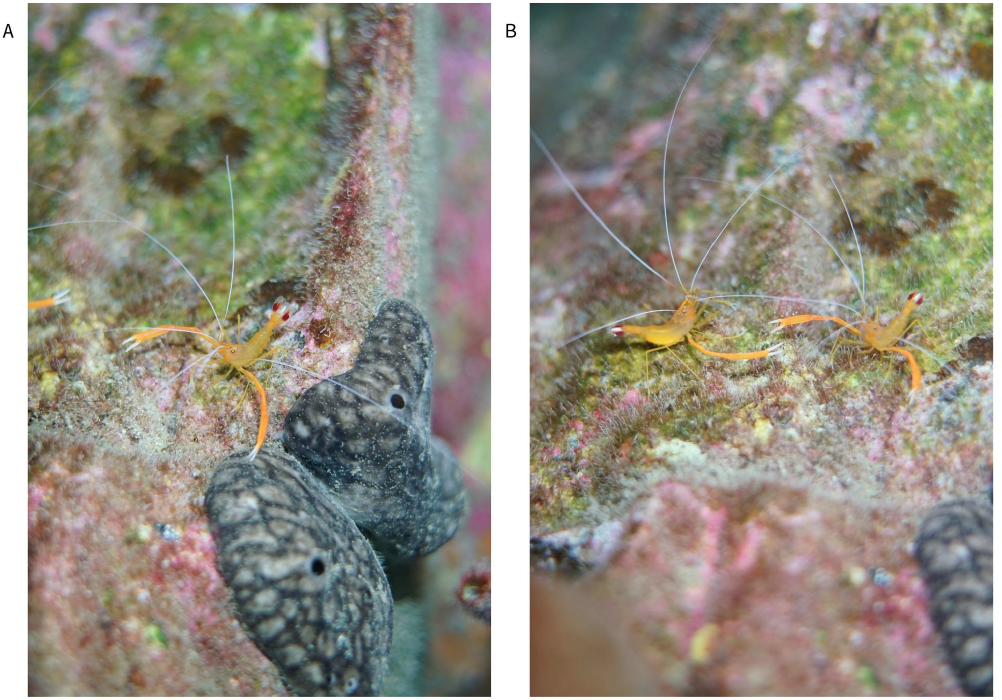
Stenopus spinosus Risso, 1827 observed in situ at Jaca, Tenerife, Canary Islands (28°07′00″N, 16°27′45″W), on 1 November 2024 during a night dive at 1:15 AM WET. Single individual emerging from rocky crevice. Two individuals observed in close proximity within a shared microhabitat. Stenopus spinosus is known to display semi-social behavior, often inhabiting narrow crevices in pairs or small groups. The species also engages in facultative cleaning behavior, and is capable of removing ectoparasites from larger reef fish. Photographs by M. Bommerer. Associated iNaturalist record: https://www.inaturalist.org/observations/250003870

### 2.3 Morphometric Measurements

Morphological measurements were obtained using digital calipers with a precision of 0.1 mm. Carapace length (CL) was measured along the dorsal midline from the postorbital margin (just posterior to the eye orbit) to the posterior edge of the carapace, excluding the rostrum. Rostrum length (RL) was measured from the base of the rostrum to its distal tip along the dorsal axis. Right cheliped length was measured as the straight-line distance from the point of articulation with the carapace to the tip of the fixed finger. Right chela length was measured as the linear distance from the proximal articulation of the propodus with the carpus to the tip of the fixed finger. All measurements conform to recognized decapod morphometric procedures as documented in the taxonomic literature

### 2.4 Field Documentation and Metadata Archiving

Two individuals of Stenopus spinosus were observed and photographed during separate night dives off Tenerife. Each observation was submitted to the iNaturalist platform, and associated metadata—including locality, date, and photographic documentation—were archived for reference. Observation details are summarized in Table 1.

**Table 1.**
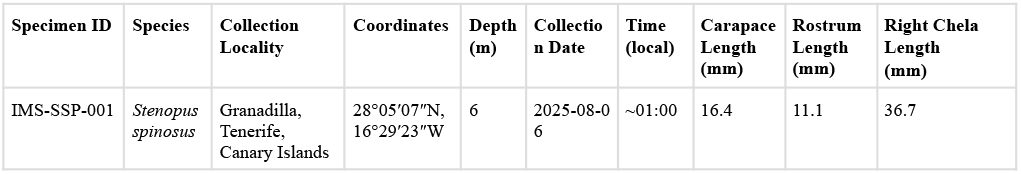

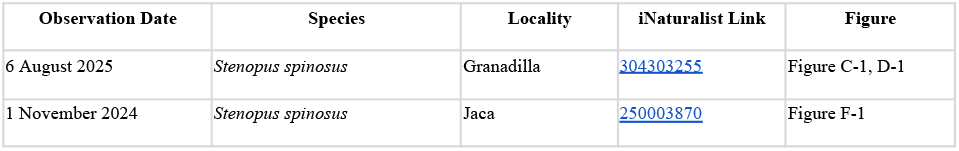

## 3. Taxonomy

The examined specimen is identified as:

- ***Stenopus spinosus*** Risso, 1827
- Family: *Stenopodidae*
- Order: *Decapoda*
- Class: *Malacostraca*

The full classification is provided below, based on DecaNet and the World Register of Marine Species (WoRMS):

Kingdom ***Animalia*** Linnaeus, 1758

Phylum ***Arthropoda*** Latreille, 1829

Subphylum ***Crustacea*** Brünnich, 1772

Superclass ***Multicrustacea*** Regier et al., 2010

Class ***Malacostraca*** Latreille, 1802

Subclass ***Eumalacostraca*** Grobben, 1892

Superorder ***Eucarida*** Calman, 1904

Order ***Decapoda*** Latreille, 1802

Suborder ***Pleocyemata*** Burkenroad, 1963

Infraorder ***Stenopodidea*** Claus, 1872

Family ***Stenopodidae*** Claus, 1872

Genus ***Stenopus*** Latreille, 1819

Species ***Stenopus spinosus*** Risso, 1827

Stenopus spinosus is the type species of the genus *Stenopus* and is currently regarded as an accepted taxon in WoRMS (AphiaID: 107721). The genus is morphologically and ecologically distinct among stenopodideans, and S. spinosus is one of the few eastern Atlantic representatives of the family.

## 4. Results

A single adult specimen of *Stenopus spinosus* Risso, 1827 was collected on 6 August 2025 from a rocky overhead environment at 6 meters depth off Granadilla, Tenerife, Canary Islands. The specimen was photographed in dorsal view under controlled lighting conditions in a laboratory setting. Two high-resolution images (Figures C–1 and D–1) were produced to illustrate the general morphology.

### Morphometric Data

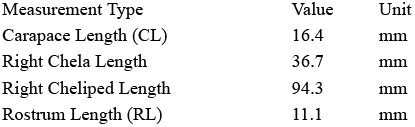

All measurements were taken along a straight axis using a calibrated ruler and digital calipers, following standard decapod morphological protocols.

- Carapace length (CL) was measured from the postorbital margin to the posterior edge of the carapace, excluding the rostrum.
- Rostrum length (RL) was recorded from the base of the rostrum to its tip along the dorsal midline.
- Right chela length refers to the straight-line distance from the articulation of the propodus with the carpus to the tip of the fixed finger (inclusive of the entire chela).
- Right cheliped length refers to the maximum straight-line distance from the point of articulation with the carapace to the distal tip of the chela (i.e., including the merus, carpus, propodus, and fixed finger).

The preserved specimen displays diagnostic morphological traits consistent with Stenopus spinosus, including a short triangular rostrum, elongate and spiny chelipeds, and distinctly banded antennae. The live coloration was a vivid reddish-orange, which faded slightly after preservation. No external parasites or deformities were observed.

### Comparative Observation

A separate in situ observation of *S. spinosus* (not part of the examined specimen) was recorded at 01:15 AM WET on 1 November 2024 by a night dive at Jaca, Tenerife (Latitude 28°07′00″ N, Longitude 16°27′45″ W). This underwater image provides ecological context and illustrates natural behavior and posture, complementing the lab-based morphological documentation.

## 5. Discussion

The genus *Stenopus* has historically been placed within the family *Stenopodidae*, yet recent molecular analyses challenge this traditional classification. A phylogenetic study by Goy et al. (2016), based on nuclear and mitochondrial gene sequences, revealed the non-monophyly of both the *Stenopodidae* and *Spongicolidae*, as well as most of their constituent genera. Despite this, *Stenopus spinosus* remains supported as a genetically coherent species within *Stenopus*, consistent with its distinct morphology.

The present record provides regionally significant morphological data for *S. spinosus* from the Canary Islands—a locality that remains underrepresented in molecular and ecological studies of *Stenopodidea*. The combination of standardized imagery, detailed morphometric measurements, and in situ ecological observations helps fill this gap and may serve as a baseline reference for future integrative taxonomic research. In particular, photographic vouchers such as those presented here—especially if paired with tissue material—can contribute to future phylogenetic assessments. These data will be useful for clarifying the evolutionary structure of stenopodid lineages.

This documentation also expands current knowledge of the species’ habitat preferences and behavior, including its tendency to shelter in crevices and its apparent capacity for semi-social or facultative cleaning interactions with reef-associated fishes. Continued fieldwork and molecular sampling across the Macaronesian region and adjacent Atlantic coasts are warranted, particularly in view of recent revisions to the evolutionary framework of *Stenopodidea*.

